# Benchmark Bias and Conformational Dynamics in Allosteric Site Prediction

**DOI:** 10.64898/2026.05.22.727284

**Authors:** Victor Pryakhin, Malika Smail-Tabbone, Yasaman Karami

## Abstract

Allosteric site prediction plays a critical role in modern drug discovery, offering opportunities to target regulatory regions with high specificity. However, most existing computational approaches rely on static protein structures and pocket detection tools such as *fpocket*, thereby overlooking conformational dynamics essential for allosteric regulation. Here, we present AlloDyn, a framework that integrates static pocket descriptors with dynamic features derived from both all-atom molecular dynamics (MD) simulations of apo-state proteins and AlphaFlow-generated conformational ensembles. By capturing structural flexibility, solvent accessibility, and residue–residue communication patterns at the pocket level, our approach enables a dynamic-aware representation of candidate allosteric sites. Importantly, we identify a systematic bias in current benchmarking practices, showing that applying *fpocket* to holo structures without removing bound allosteric modulators introduces data leakage and leads to artificially inflated performance estimates. When evaluated on properly preprocessed datasets, dynamic feature augmentation significantly improves prediction performance over static baselines. Furthermore, we demonstrate that AlphaFlow-generated ensembles achieve performance comparable to MD-derived features at a fraction of the computational cost, providing a scalable alternative for conformational sampling. Benchmarking on the D24 dataset shows that AlloDyn achieves the best balance between precision and recall, yielding the highest F1 score and MCC among evaluated methods. We show that current benchmarks overestimate performance due to data leakage, and that incorporating dynamics is key to accurate and scalable allosteric site prediction.

## Introduction

Allostery is a fundamental regulatory mechanism in which the binding of a molecule (*effector* or *modulator*) at one site of a protein (the *allosteric site*) induces functional changes at a distant site, often the *active site*. This long-range intramolecular communication enables proteins to modulate their activity in response to cellular signals. Allosteric regulation is widespread across biological systems and plays a central role in controlling enzymatic catalysis, metabolic pathways, and signal transduction [1]. Importantly, effectors can act as activators or inhibitors, finely tuning protein function without directly interacting with the substrate-binding site [2].

Allosteric sites often lie outside conserved active regions, making them attractive yet challenging targets for drug design. Unlike orthosteric drugs, allosteric modulators can offer enhanced specificity and reduced toxicity [3, 4]. However, the structural diversity of allosteric mechanisms across protein families has long complicated efforts to systematically identify allosteric sites.

The systematic collection of experimental annotations in recent years has greatly improved the accessibility of curated data on allosteric proteins. The Allosteric Database (ASD) [5] is the most comprehensive, with derived subsets such as ASBench [6] and CASBench [7] providing filtered, benchmarking-oriented collections. AlloBench [8] was recently introduced to facilitate standardized evaluation of predictive models. The growing volume and accessibility of annotated allosteric data have enabled the development of data-driven computational approaches for allosteric site prediction, offering scalable and generalizable alternatives to experimental identification.

Computational methods for allosteric site prediction can be broadly divided into two categories. *Residue-level* methods aim to identify individual allosteric residues within a protein, typically by combining structural, evolutionary, and dynamic features at the per-residue level, as in AR-Pred [9], which couples dynamics features using elastic network models with evolutionary information, NACEN [10], which exploits residue contact energy networks, and more recently a nanoenvironment descriptor-based approach [11]. The advent of protein language models (PLMs) has further enabled *sequence-based* residue prediction without requiring a three-dimensional structure, as demonstrated by AlloFusion [12], which leverages PLM embeddings, and Kannan et al. [13], which uses PLM attention maps to identify allosteric residues directly from sequence. *Pocket (or site)-level* methods, by contrast, operate on cavities detected on the protein surface and classify them as allosteric or non-allosteric, offering a more directly actionable output for drug design. While residue-level approaches provide finer mechanistic insight, pocket-level methods are better suited for identifying druggable sites and are the focus of the present work.

Pocket-level approaches have diversified over the past decade, yet most machine learning-based models still rely on static protein conformations. Many such methods use geometry-based pocket detection tools such as *fpocket* [14] to derive physicochemical features and employ classifiers, as implemented in *Allosite* [15] and the *PASSer* family [16–18]. Other recent strategies extend these approaches by incorporating complementary structural descriptors and advanced feature selection, as in *MEF-AlloSite* [19]. Beyond static representations, several methods rely on coarse-grained dynamic analysis. *AllositePro* [20] uses normal mode analysis based on an elastic network model to evaluate protein dynamic changes upon allosteric ligand binding; *AlloPred* [21] applies normal mode perturbation analysis to capture flexibility changes, and *ESSA* (Essential Site Scanning Analysis) [22] identifies residues whose perturbation strongly alters the dispersion of global motions within elastic network-based normal modes. A related perturbation-based approach, *APOP* [23], evaluates the allosteric potential of fpocket-detected pockets by stiffening residue–residue interactions within each cavity in a Gaussian network model and ranking them based on induced shifts in global mode frequencies combined with hydrophobicity descriptors. More recently, *AllosES* [24] expanded this concept by integrating transfer entropy–based coupling derived from Gaussian network models, followed by *AlloEF* [25] and *ZHMolEReP* [26], two residue-level methods that combine dynamic and energetic features (transfer entropy and energetic frustration for the former, perturbation response scanning and free energy approximation for the latter), both requiring active site information as input.

While the coarse-grained dynamic analyses described above go beyond purely static representations, they rely on simplified physical models and a single input structure, which may not fully capture the conformational heterogeneity central to allosteric regulation. Molecular dynamics (MD) simulations offer a principled route to capture such dynamics, sampling the conformational space of a protein at atomic resolution over time. However, the computational cost of all-atom MD simulations remains substantial, particularly at the scale of large benchmark datasets. The recent emergence of generative models for conformational sampling, such as AlphaFlow [27], offers a promising and scalable alternative. More broadly, the growing interest in dynamics-aware machine learning, including methods that incorporate conformational flexibility into binding interface prediction [28], underscores the potential of integrating protein dynamics into allosteric site modelling.

In this study, we present AlloDyn, which is, to the best of our knowledge, the first dynamic-aware method for allosteric site prediction that integrates static *fpocket* descriptors with dynamic features derived from full-atom MD simulations of apo (modulator-free) protein structures, allowing direct comparison with a static-only baseline. We further evaluate AlphaFlow [27], a diffusion-based generative model that constructs conformational ensembles directly from protein sequences, as a computationally tractable alternative to MD simulations. In addition, we identify and characterize a systematic bias introduced when *fpocket* is applied to holo structures without prior removal of the allosteric modulator, leading to data leakage and artificially inflated performance estimates. AlloDyn is evaluated against several state-of-the-art allosteric site prediction methods on the D24 benchmark set [20], demonstrating competitive and consistent classification performance.

## Materials and Methods

### Allosteric Dataset Construction and Annotation

The dataset used in this study was prepared following a pipeline inspired by PASSerRank [18], which was developed to curate a high-quality, structurally consistent, and sequence-diverse collection of protein–modulator complexes for allosteric site prediction. Protein structures and annotations were obtained from the 2023 release of the Allosteric Database (ASD2023) [5], which contains over 2,400 experimentally validated protein–modulator entries. To ensure structural reliability, we retained only X-ray structures with resolution better than 3.0 Å, and excluded entries with incomplete allosteric sites or ambiguous modulator annotations. Pocket identification was performed using *fpocket* [14], a widely used geometry-based method. We explored two preprocessing strategies prior to pocket detection. In the first, *fpocket* was applied directly to the full structure using a flag to restrict detection to the chain annotated as allosteric in the ASD, without any prior cleaning. In the second, solvent molecules and ions were removed, the allosteric modulator was extracted from the chain, and *fpocket* was then applied to the cleaned, isolated chain. For each protein, the geometric centers (*centroids*) of the predicted pockets and the modulator were computed. The pocket with the shortest centroid-to-centroid distance to the modulator (*pocket-modulator distance*) was labeled as allosteric, and all others as non-allosteric. This modulator-based labeling strategy was adopted as residue-level allosteric site annotations are not consistently available across all entries in ASD. If the minimum distance exceeded 10 Å (*default threshold*), the entry was excluded, as such cases typically reflect failures in pocket detection [18]. Redundancy reduction was applied to obtain a sequence-diverse dataset, using pairwise sequence identity filtering with a 30% threshold. After detection, labeling, and filtering the final dataset consisted of 353 proteins, comprising 8,847 pockets in total, of which 353 were labeled as allosteric and 8,494 as non-allosteric (Figure 1-(1)).

**Figure 1:**
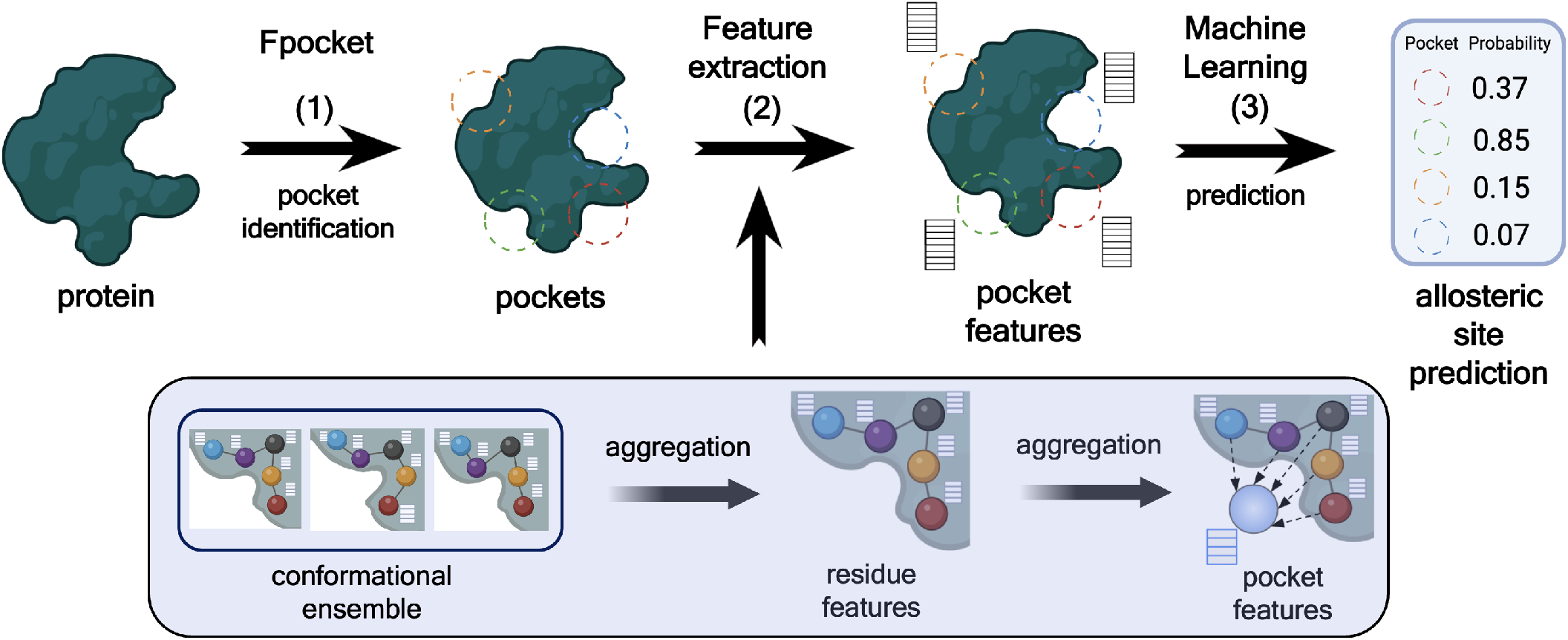
Overview of the AlloDyn pipeline. (1) *fpocket* detects pockets on the input protein structure. (2) Static features come directly from *fpocket* descriptors; dynamic features come from conformational ensembles generated by MD simulations or AlphaFlow, aggregated across the ensemble into residue-level features and then into pocket-level features. (3) XGBoost combines both feature sets and outputs an allosteric probability score for each pocket.

### Generation of Conformational Ensembles

#### Molecular Dynamics simulations

For each of the 353 protein chains, we performed three replicates of 500-ns MD simulations, starting from their respective experimental structures. All bound ligands were removed, while ions were retained. Missing residues in regions with continuous gaps were modeled using MODELLER [29]. MD simulations were performed following the protocol described in [30], with identical simulation settings. Simulations were performed using GROMACS 2024.2 [31] with the CHARMM36m force field [32]. Systems were neutralized and adjusted to physiological ionic strength (150 mM NaCl) by adding Na^+^ and Cl^−^ ions. Each system was solvated in a dodecahedral box of explicit TIP3P water model with a minimum buffer distance of 12 Å from the solute. Hydrogen atoms were added, and histidine protonation states were assigned using Reduce [33]. Energy minimization was carried out using the steepest descent algorithm for 5000 steps to remove steric clashes. This was followed by a multi-step equilibration phase in the NPT ensemble at 310 K, during which positional restraints were applied to protein and lipid heavy atoms and gradually released over 0.375 ns. Pressure was maintained at 1 bar using the Berendsen barostat during equilibration [34]. Production simulations were performed in the NPT ensemble with a time step of 2 fs. For each system, three independent replicates of 500 ns were generated using different initial velocities. Temperature was maintained at 310 K using the V-rescale thermostat [35], and pressure was controlled at 1 atm using the C-rescale barostat under isotropic conditions [36]. Covalent bonds involving hydrogen atoms were constrained using the LINCS algorithm [37], and long-range electrostatic interactions were treated with the Particle Mesh Ewald method [38]. Coordinates were recorded every 100 ps. For downstream analysis,the three MD replicates were combined into a single concatenated trajectory for each protein, which served as the basis for dynamic feature extraction.

#### Generative models

As an alternative to classical MD simulations, we explored the use of AlphaFlow [27], a diffusion-based generative model that produces conformational ensembles directly from protein sequences. We employed its distilled version, which offers a favorable tradeoff between accuracy and computational efficiency. For consistency with MD-derived ensembles, we used the same curated protein sequences as input to AlphaFlow. For each protein entry, 25 conformations were generated. To ensure consistent atom and residue indexing, the sequences were aligned to their corresponding experimental structures prior to feature extraction.

### Static and Dynamic Features Construction

#### Fpocket features

Each detected pocket was described using 19 descriptors computed by *fpocket*. These features capture geometric, physicochemical, and topological properties of the pocket, and include descriptors such as volume, solvent-accessible surface area (SASA), polarity, hydrophobicity, charge, and alpha-sphere statistics. Although one of these descriptors encodes residue flexibility via B-factors, all fpocket descriptors are derived from a single static structure rather than from conformational ensembles, and are referred to as *static* or *fpocket* features throughout this work.

#### Dynamic features

We computed dynamic descriptors from conformational ensembles to capture various aspects of pocket dynamics (Figure 1-(2)). Pockets were defined on the experimental structures using *fpocket* and represented as fixed sets of residues. Dynamic descriptors were computed by tracking these residues across conformational ensembles at the pocket level. Dynamic features were organised into three groups as described below (full list of features is available in Supplementary Table S1).

#### Compactness and shape features

All features in this group were computed from the C*α* atom coordinates of pocket residues using MDAnalysis [39]. For each frame, the pocket centroid was computed as 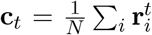 over the *N* C*α* atoms of the pocket. Based on this definition, we extracted nine descriptors capturing pocket geometry and variability across the ensemble. These include the standard deviation of the centroid position, as well as the mean and standard deviation of the minimum, average, and maximum C*α*-to-centroid distances. In addition, mean pairwise inter-residue distances were computed per frame and summarised by their mean and standard deviation across the ensemble. Pocket compactness was quantified using the root mean square radius, 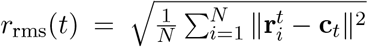, computed per frame and summarised by its mean and standard deviation. Residue-level flexibility was captured through per-residue RMSF values, aggregated as mean and standard deviation across pocket residues. Finally, pocket shape was characterised per frame by the eigenvalues of the 3*×*3 spatial covariance matrix of C*α* positions, 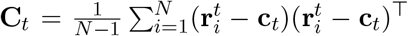. The resulting eigenvalues λ_1_ *≥* λ_2_ *≥* λ_3_ describe the principal axes of the instantaneous spatial distribution of pocket residues, and were used to derive anisotropy *A* = λ_3_*/*λ_1_ and sphericity *S* = (λ_1_λ_2_λ_3_)^1*/*3^*/*(λ_1_ + λ_2_ + λ_3_), both summarised by their mean and standard deviation over the ensemble (17 features).

#### SASA-based features

Pocket solvent-accessible surface area is computed per frame using MDTraj [40]. Both absolute (Å^2^) and relative (fraction of total protein SASA) values are recorded, with mean, standard deviation, range, and coefficient of variation computed for each, leading to a total of 8 features per pocket.

#### Dynamic coupling features

Pairwise couplings between all protein residues are quantified using four matrices. The *dynamic cross-correlation* (Pearson) measures the normalised cross-covariance of C_*α*_ positional fluctuations [41]:

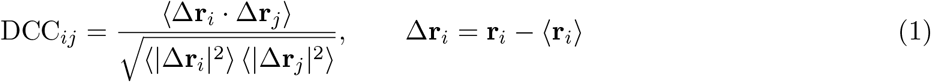

*Mutual information* (MI) and *generalised correlation* (GC) capture both linear and nonlinear couplings [42]. Under the Gaussian approximation:

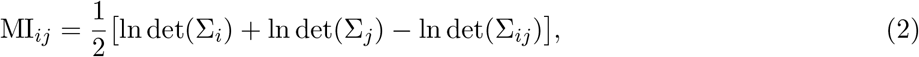

whereΣ_*i*_,Σ_*j*_ are the covariance matrices of residues *i* and *j*, andΣ_*ij*_ is their joint covariance matrix. GC [42] is derived from MI as:

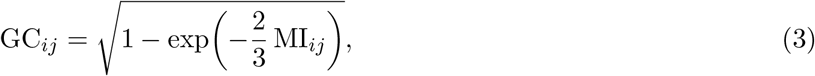

Finally, the *communication propensity* (CP) quantifies the variance of the inter-residue distance across the ensemble [43]:

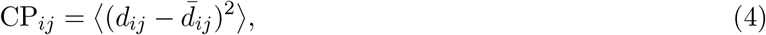

where *d*_*ij*_ is the instantaneous distance between C_*α*_ atoms of residues *i* and *j*. GC, MI, and CP were computed using ComPASS [44]. For each matrix, three sets of pocket-level statistics are derived: *intra-pocket* couplings (mean, max, and std of all residue pairs within the pocket), *global connectivity* (mean and std of couplings between pocket residues and the rest of the protein), and *inter-pocket* couplings (statistics over couplings between this pocket and every other detected pocket).

### Model Training and Evaluation

#### Classifier and hyperparameter optimization

We trained gradient-boosted decision tree models using XGBoost [45], selected for its speed, scalability, and effectiveness on imbalanced classification tasks (Figure 1-(3)). To prevent data leakage and ensure generalization across protein structures, we employed a group-aware splitting strategy: both the train/test split (80/20) and the 5-fold cross-validation for hyperparameter tuning were performed such that all pockets from the same protein were consistently assigned to a single partition. Hyperparameters were optimized via grid search, with model selection based on the average F1 score across cross-validation folds. To address the substantial class imbalance between allosteric and non-allosteric pockets, the `scale_pos_weight` parameter, which controls the weight assigned to the positive class, was included in the hyperparameter search space. The best-performing configuration was used to retrain the model on the full training set and evaluated on the independent test set.

#### Evaluation of dynamic feature contribution

To obtain robust performance estimates, the full training and evaluation pipeline was repeated across 50 random group-based train/test splits. This procedure was applied to the static baseline model, as well as to models augmented with dynamic features from MD simulations or AlphaFlow-generated conformational ensembles. To ensure a fair comparison between MD- and AlphaFlow-derived features, only entries with fewer than 862 residues were retained, as this represented the upper length limit for AlphaFlow, resulting in 339 entries (7,779 pockets, non-allosteric-to-allosteric ratio *∼*22:1). Performance was evaluated primarily using the F1 score, given the strong class imbalance; precision, recall, MCC, AUC-ROC, and AUC-PR were additionally computed.

#### Benchmarking against state-of-the-art methods

To compare our approach with existing methods, we used the D24 benchmark set from AllositePro [20]. Entries sharing high sequence identity with D24 proteins were removed from our training dataset using pairwise sequence identity filtering at a 30% threshold, yielding a filtered training set of 293 entries (6,704 pockets, class ratio *∼*22:1). The final model was trained on this filtered dataset using 5-fold group-aware cross-validation to select optimal hyperparameters, followed by retraining on the full filtered set. This model was then evaluated on D24 alongside PASSer (ensemble [16], AutoML [17], and ranking-based [18] variants), AllosES [24], and APOP [23]. D24 structures were processed using the same pipeline applied to our main dataset: ligands, solvent molecules, and ions were removed, and for each entry, only the single functional chain annotated as allosteric in the AllositePro set was retained (Supplementary Table S2). The pocket closest to the bound modulator was labeled as the allosteric site, resulting in exactly one positive entry per structure. For all methods, precision, recall, F1-score, MCC, AUC-ROC, and AUC-PR were computed; for methods providing only ranking scores (APOP, PASSer-ranking), all positively ranked pockets were treated as predicted positives prior to metric computation.

## Results

### Modulator Presence Introduces a Systematic Bias in Fpocket Descriptors

When *fpocket* is applied directly to a holo structure without prior removal of the allosteric modulator, the resulting pocket descriptors can differ substantially from those obtained after pre-cleaning. We refer to these two strategies as *biased* (no pre-cleaning) and *unbiased* (pre-cleaned structure) throughout this work. We identified two sources of inconsistency introduced by the biased approach. First, in cases where the allosteric modulator is a modified amino acid, *fpocket* may incorporate it into the pocket definition or even construct a pocket around it, directly biasing the pocket geometry and composition. Second, and more systematically, we observed that while *fpocket* does not consider the modulator during pocket *detection*, it appears to include it during *feature calculation*. In particular, SASA values are computed over the full structure including the modulator, which artificially reduces the solvent-exposed area of the allosteric pocket in the biased case. Since the *fpocket* score and druggability score are derived from regression formulas that incorporate SASA, this propagates into a systematic upward shift of these scores for allosteric pockets in the biased dataset. This effect is clearly visible in the score distributions (Figure 2A): while non-allosteric pockets show nearly identical distributions between the biased and unbiased configurations, allosteric pockets exhibit a pronounced rightward shift in *fpocket* score and druggability score under the biased configuration. Manual inspection of individual entries (Figure 2B-C, Supplementary Table S3) confirmed that the primary differences between biased and unbiased descriptors are concentrated in *fpocket* score, druggability score, and SASA-related features, while geometric descriptors such as volume, alpha-sphere statistics, and flexibility remained largely unchanged.

**Figure 2:**
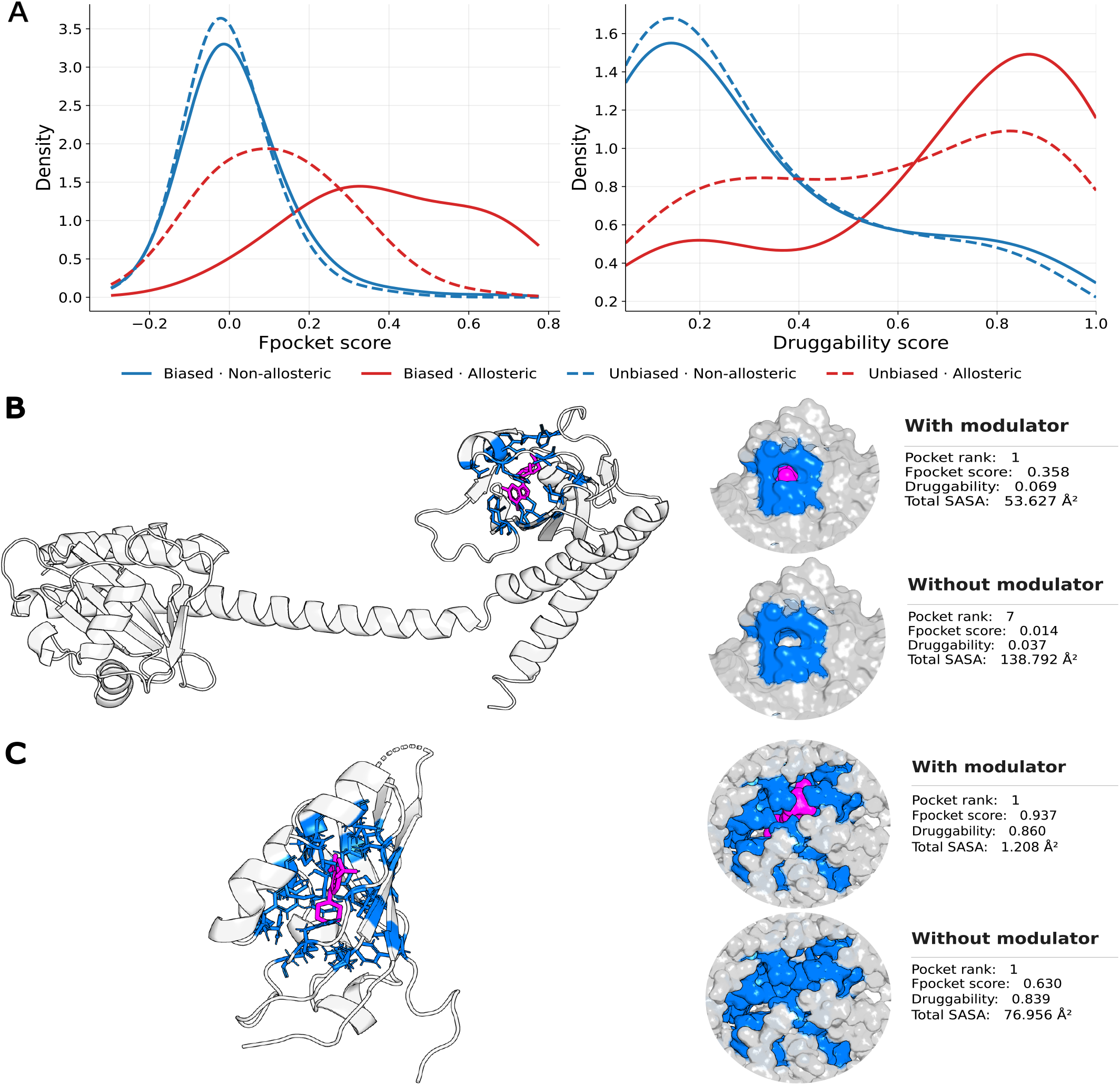
Comparison of fpocket descriptors between biased and unbiased configurations. **(A)** Kernel density estimates of fpocket score and druggability score across all detected pockets, split by allosteric and non-allosteric labels. **(B)** 1MC0 and **(C)** 3F1O allosteric binding sites shown in biased (modulator present) and unbiased (modulator removed) configurations. Left: full protein structure (cartoon) with allosteric residues (marine sticks) and modulator (magenta sticks). Right: zoomed surface views of the allosteric pocket (marine) with (top) and without (bottom) the modulator (magenta surface). Removal of the modulator affects the *fpocket* score (and potentially the pocket rank), druggability score, and total SASA, illustrating the systematic bias introduced when *fpocket* is applied to holo structures.

The impact of this data leakage is directly reflected in model performance (Table 1): models trained on the biased dataset achieve substantially higher F1 scores than those trained on the unbiased dataset.

**Table 1:** Model performance on biased and unbiased datasets, averaged over 50 random seeds. For each dataset, results are reported for the *fpocket*-only baseline and AlloDyn models augmented with MD- or AlphaFlow-derived (AF) dynamic features. Values are reported as mean *±* standard deviation of test set performance across the 50 splits.

| Source | Model | Precision | Recall | F1 | MCC | AUC ROC | AUC PR |
| --- | --- | --- | --- | --- | --- | --- | --- |
| Biased | Baseline | $0.57 \pm 0.05$ | $0.65 \pm 0.09$ | $0.60 \pm 0.05$ | $0.59 \pm 0.05$ | $0.94 \pm 0.02$ | $0.61 \pm 0.06$ |
| | AlloDyn (Fpocket+MD) | $0.54 \pm 0.03$ | $0.69 \pm 0.09$ | $0.61 \pm 0.04$ | $0.59 \pm 0.05$ | $0.95 \pm 0.01$ | $0.59 \pm 0.04$ |
| | AlloDyn (Fpocket+AF) | $0.56 \pm 0.05$ | $0.68 \pm 0.08$ | $0.61 \pm 0.05$ | $0.60 \pm 0.05$ | $0.95 \pm 0.02$ | $0.62 \pm 0.05$ |
| Unbiased | Baseline | $0.46 \pm 0.06$ | $0.41 \pm 0.06$ | $0.43 \pm 0.05$ | $0.41 \pm 0.05$ | $0.86 \pm 0.03$ | $0.40 \pm 0.06$ |
| | AlloDyn (Fpocket+MD) | $0.48 \pm 0.06$ | $0.42 \pm 0.05$ | $0.45 \pm 0.05$ | $0.43 \pm 0.05$ | $0.86 \pm 0.02$ | $0.41 \pm 0.06$ |
| | AlloDyn (Fpocket+AF) | $0.48 \pm 0.06$ | $0.42 \pm 0.06$ | $0.45 \pm 0.05$ | $0.43 \pm 0.05$ | $0.87 \pm 0.02$ | $0.40 \pm 0.06$ |

However, this apparent performance gain is artifactual: feature importance analysis (Supplementary Figure S1) confirms that the *fpocket* score is the dominant predictor in the biased setting, disproportionately driving classification due to its artificially inflated values for allosteric pockets. This inflated performance does not reflect the model’s ability to identify genuine allosteric features, and cannot be used as a reliable estimate of predictive performance.

Taken together, these findings indicate that applying *fpocket* to uncleaned holo structures introduces a data leakage: allosteric pockets receive artificially inflated scores due to the presence of the modulator, making them easier to classify for reasons unrelated to genuine structural features. The unbiased preprocessing strategy, in which the modulator is removed prior to pocket detection, eliminates this artifact and provides a more reliable basis for model training and evaluation.

### Dynamic Feature Augmentation Improves Allosteric Site Prediction

To evaluate the contribution of dynamic features to allosteric site prediction, we compared models trained on *fpocket* descriptors alone (baseline) to those augmented with dynamic features derived from MD simulations or AlphaFlow-generated conformational ensembles, using the unbiased dataset. Model training and evaluation were repeated across 50 random group-based train/test splits. Statistical significance of performance differences was assessed using two-tailed paired *t*-tests on matched splits, after confirming normality of all metric distributions with the Shapiro–Wilk test (*p >* 0.05); effect sizes were quantified using Cohen’s *d* (more details in Supplementary Table S4).

Both AlloDyn variants significantly outperformed the static baseline across multiple metrics (Table 1). For F1 and MCC, improvements were statistically significant (^*\*\**^*p <* 0.01) with small effect sizes (|*d*| *≈* 0.4). For AUC-PR, the MD-augmented model reached a medium effect size (|*d*| = 0.56, ^*\*\*\**^*p <* 0.001). Models trained on MD- and AlphaFlow-derived features performed comparably across all metrics, with no statistically significant difference between the two conformational sources for F1 and MCC (*p >* 0.98, |*d*| *≈* 0.00). This indicates that AlphaFlow-generated ensembles can serve as a computationally efficient alternative to MD simulations within the current classification framework.

### Comparison with State-of-the-Art Methods

AlloDyn was evaluated on the D24 benchmark set against PASSer (ensemble [16], AutoML [17], and ranking-based [18] variants), AllosES [24], and APOP [23]. For this evaluation, the AlloDyn model was trained using AlphaFlow-generated conformational ensembles as the source of dynamic features. All methods were evaluated using the same set of classification metrics (Table 2). For methods providing only ranking scores (APOP, PASSer-ranking), all positively ranked pockets were treated as predicted positives when computing classification metrics. The number of evaluated pockets (*N*) varies across methods. AlloDyn and PASSer operate on the same pocket set (*N* = 645). The small discrepancy for APOP (*N* = 641) arises from its mandatory removal of non-standard amino acids during preprocessing, which introduces gaps in the sequence and alters the alpha-sphere clustering step in *fpocket*, resulting in slightly different pocket definitions for a small number of entries. For AllosES, the use of an older version of *fpocket* in combination with non-standard amino acid removal led to a substantially different pocket set (*N* = 305).

**Table 2:** Performance comparison of allosteric site prediction methods on the D24 benchmark. Metrics are reported at the pocket level. *N* denotes the number of evaluated pockets. Best values per column are shown in bold.

| Method | Precision | Recall | F1 | MCC | AUC-ROC | AUC-PR | N |
| --- | --- | --- | --- | --- | --- | --- | --- |
| AlloDyn | 0.52 | 0.50 | **0.51** | **0.49** | **0.93** | 0.44 | 645 |
| APOP | 0.09 | **0.95** | 0.17 | 0.23 | **0.93** | 0.52 | 641 |
| AllosES | 0.24 | 0.79 | 0.37 | 0.37 | 0.86 | **0.54** | 305 |
| AllosES* | **0.60** | 0.27 | 0.38 | 0.39 | 0.90 | 0.41 | 645 |
| PASSer-automl | 0.50 | 0.27 | 0.35 | 0.35 | 0.91 | 0.32 | 645 |
| PASSer-ensemble | 0.45 | 0.23 | 0.30 | 0.31 | 0.86 | 0.38 | 645 |
| PASSer-rank | 0.21 | 0.14 | 0.17 | 0.15 | 0.84 | 0.28 | 645 |

\*Re-evaluated with *fpocket* v.4 to match the pocket set used by all other methods ( $N = 645$ ).

Figure 3 illustrates representative prediction outcomes on the D24 benchmark. Panel **A** shows a prediction example for 4OYA_A, human soluble adenylate cyclase, demonstrating the discriminative power of AlloDyn: while AllosES and PASSer variants assign the correct allosteric pocket a lower score and instead prioritize a distant, non-allosteric cavity, AlloDyn correctly identifies it as the top prediction. Panels **B–C** illustrate the fundamental limitation imposed by the pocket detection step: for 5J94_A and 4NHV_A, no *fpocket*-detected pocket falls within the 10 Å labelling threshold of the allosteric modulator, meaning all pockets were labelled as non-allosteric and the correct site could not be recovered by any method. Detailed predictions for all D24 entries across all compared methods are provided in Supplementary Figures S2–S4.

**Figure 3:**
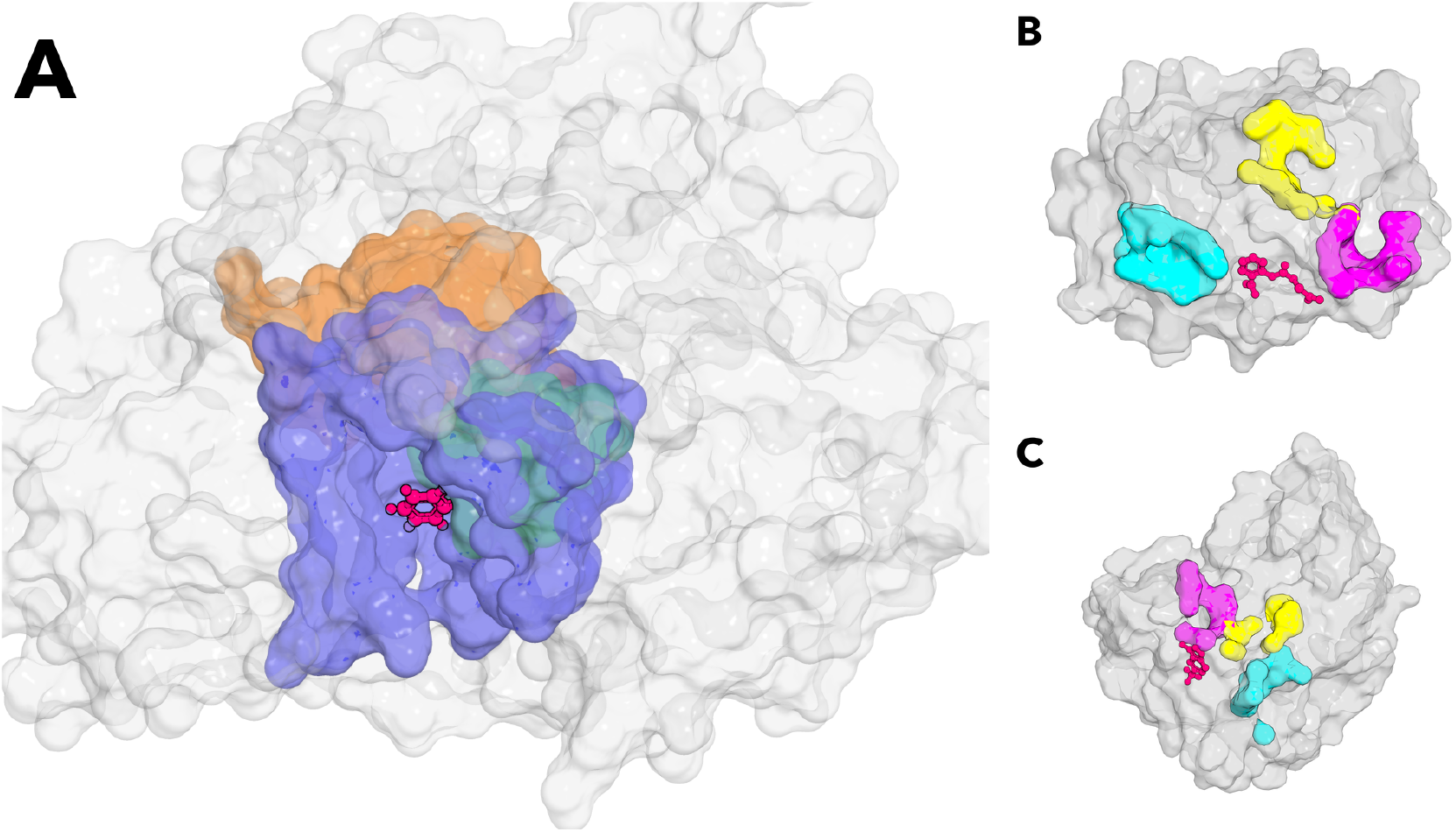
Representative allosteric pocket predictions on the D24 benchmark. **(A)** Prediction comparison for 4OYA_A. The correct allosteric pocket is ranked first by AlloDyn (green, buried pocket, 2.5Å from modulator) and APOP (blue, larger binding-site representation). The difference in pocket extent between the two methods arises from APOP’s removal of modified residues during preprocessing. AllosES and PASSer variants assign a lower rank to the correct allosteric pocket, instead prioritizing a distant pocket (orange, 15.3 Å from the modulator) as their top prediction. Modulator is shown in pink (sticks). **(B–C)** Failure cases for 5J94_A and 4NHV_A, respectively. No *fpocket*-detected pocket falls within the 10 Å labelling threshold of the allosteric modulator (pink sticks), showing that prediction performance is fundamentally constrained by the pocket detection step in these cases. The three closest detected pockets are shown: the nearest (magenta), second nearest (cyan), and third nearest (yellow). Protein surface is shown in light gray.

Among all evaluated methods, AlloDyn achieves the best balance between precision and recall, yielding the highest F1 score (0.51) and MCC (0.49, Table 2). APOP attains the highest recall (0.95) together with a competitive AUC-PR (0.52), but at the cost of very low precision (0.09), indicating that a large fraction of detected pockets are predicted as allosteric, leading to numerous false positives. Similarly, AllosES achieves the highest AUC-PR (0.54), suggesting strong ranking performance across thresholds, but its lower precision results in reduced F1 and MCC scores compared to AlloDyn. PASSer variants show moderate and consistent performance across metrics, with PASSer-AutoML performing best within the family. These results indicate that while APOP and AllosES are effective at assigning elevated scores to allosteric pockets, they tend to overpredict positive sites. In contrast, AlloDyn provides a more balanced tradeoff between sensitivity and specificity, leading to more reliable threshold-based classification performance. In practical applications, where experimental validation of predicted allosteric sites is costly, reducing false positives while maintaining competitive recall may represent a more useful operating regime.

## Discussion

This study investigated whether dynamic features derived from conformational ensembles can improve allosteric site prediction beyond what is achievable with static *fpocket* descriptors alone. Using a curated dataset of annotated allosteric and non-allosteric pockets, we trained and evaluated XGBoost classifiers on static features, conformational ensemble-derived dynamic features (MD or AlphaFlow-generated). Our results show that dynamic feature augmentation significantly improves classification performance when structures are properly preprocessed, while also revealing several methodological bottlenecks that limit the utility of dynamic descriptors in the current framework.

Pocket detection constitutes the first and most critical step in our pipeline, and its accuracy directly determines the upper bound of downstream prediction performance. When *fpocket* is applied to holo structures without prior removal of the allosteric modulator, SASA-based descriptors are computed over the full structure including the modulator, artificially inflating the *fpocket* score and druggability score for allosteric pockets. This constitutes a data leakage that leads to overoptimistic performance estimates. Removing the modulator prior to pocket detection eliminates this artifact and provides a more reliable basis for model training. Beyond preprocessing, *fpocket* may simply fail to detect the correct allosteric pocket in some cases, as illustrated by the two D24 entries (5J94_A and 4NHV_A) where no detected pocket fell within the labelling threshold of the modulator. In such cases, no downstream method can recover the correct site, highlighting the fundamental dependency of our approach on pocket detection quality.

Despite the dominance of static descriptors, dynamic feature augmentation led to a statistically significant improvement in F1 score on the unbiased dataset. Notably, some static *fpocket* descriptors (in particular the *fpocket* score and druggability score) remained among the top predictors across all settings, reflecting their strong discriminative power even in the absence of dynamic information. These results suggest that dynamic features are most informative when the allosteric pocket is accurately identified, highlighting the dependency of dynamic descriptors on the quality of the initial pocket definition.

Dynamic features were computed by anchoring to the set of pocket-forming residues identified in the experimental structure. While this provides a consistent reference across conformations, it is an approximation: the true pocket boundary may shift during simulation, and transient or cryptic regions may not be captured by residues defined on a single static structure. More sophisticated pocket tracking strategies, such as ensemble-based pocket detection or adaptive residue mapping across conformations, could improve the accuracy of dynamic descriptors. Furthermore, obtaining pocket-level features requires a double aggregation - from the conformational ensemble to per-residue features, and then from residues to the pocket level - which inevitably discards spatial and temporal resolution. More expressive architectures, such as graph neural networks with learnable aggregation, could better preserve this information and more fully exploit the rich structural information contained in conformational ensembles.

Correlation-based features capture residue-residue communication patterns that are conceptually well-suited to allosteric regulation. However, their computation requires knowledge of which residues belong to each pocket, and the current approach uses all detected pockets for inter-pocket correlation calculations due to the absence of explicit orthosteric site annotations in the dataset. This introduces noise in the inter-pocket and global connectivity features, as non-functional pockets are treated on equal footing with biologically relevant ones. Future work incorporating orthosteric site annotations would allow more targeted and interpretable correlation descriptors.

Despite known structural imperfections such as occasional atomic clashes, models trained on AlphaFlow-derived dynamic features achieved performance comparable to those using MD-derived ensembles, with no statistically significant differences. A current limitation of AlphaFlow is that it operates on single protein chains, whereas MD simulations can accommodate multimeric complexes. For proteins whose allosteric regulation depends on inter-chain interactions, this represents a meaningful constraint. Nevertheless, for monomeric systems, AlphaFlow offers a practical and computationally efficient alternative to MD for conformational sampling in this context.

Overall, our results demonstrate that integrating dynamic features into allosteric site prediction is a promising direction, but its effectiveness is conditioned on reliable pocket detection, accurate structural annotations, and sufficiently expressive feature representations. Expanding annotated datasets with orthosteric site information and developing architectures capable of learning directly from raw conformational ensembles are key directions for future work.

## Data and code availability

The source code and training datasets are publicly available on GitLab at https://gitlab.inria.fr/vpryakhi/AlloDyn.

## Acknowledgements

YK was supported by the French National Research Agency (ANR) under the France 2030 grant reference number ANR-24-RRII-0002 operated by the Inria Quadrant Program. Computations were performed using resources from the Grid’5000 and MBI computing platforms. This work was granted access to the HPC resources of IDRIS under the allocation 2025-A0180714660 granted to Y.K. made by GENCI.

