## Supplementary Information for "Benchmark Bias and Conformational Dynamics in Allosteric Site Prediction"

### Supplementary Tables

Table S1: Complete list of dynamic feature symbols used in AlloDyn. The wildcard * denotes one of {dcc, gc, mi, cp} unless stated otherwise.

| Feature | Symbol(s) |
|---|---|
| ***Structural dynamics*** | |
| Centroid position std | `centroid_std` |
| Root mean square radius | `rg_mean`, `rg_std` |
| Min Cα-to-centroid distance | `cent_min_dist_mean`, `cent_min_dist_std` |
| Avg Cα-to-centroid distance | `cent_avg_dist_mean`, `cent_avg_dist_std` |
| Max Cα-to-centroid distance | `cent_max_dist_mean`, `cent_max_dist_std` |
| Inter-residue distance | `interres_avg_dist_mean`, `interres_avg_dist_std` |
| RMSF | `rmsf_mean`, `rmsf_std` |
| Anisotropy | `anisotropy_mean`, `anisotropy_std` |
| Sphericity | `sphericity_mean`, `sphericity_std` |
| ***SASA-based features*** | |
| SASA absolute | `sasa_mean_abs`, `sasa_std_abs`, `sasa_range_abs`, `sasa_cv_abs` |
| SASA relative | `sasa_mean_rel`, `sasa_std_rel`, `sasa_range_rel`, `sasa_cv_rel` |
| ***Intra-pocket coupling features*** | |
| Number of pocket residues | `n_pocket_residues_corr` |
| Intra-pocket coupling mean | `*_intra_mean` |
| Intra-pocket coupling max | `*_intra_max` |
| Intra-pocket coupling std | `*_intra_std` |
| High coupling fraction | `gc_high_corr_frac`, `pearson_high_corr_frac` |
| Pearson intra min | `pearson_intra_min` |
| Pearson intra absolute mean | `pearson_intra_abs_mean` |
| Pearson intra absolute max | `pearson_intra_abs_max` |
| Pearson intra absolute std | `pearson_intra_abs_std` |
| Pearson intra positive fraction | `pearson_intra_pos_frac` |
| Pearson intra negative fraction | `pearson_intra_neg_frac` |
| ***Global connectivity features*** | |
| Global coupling mean | `*_global_mean` |

Continued on next page

Table S1 – continued from previous page

| **Feature** | **Symbol(s)** |
|---|---|
| Global coupling std | `*_global_std` |
| Pearson global absolute mean | `pearson_global_abs_mean` |
| Pearson global absolute std | `pearson_global_abs_std` |
| Pearson global positive fraction | `pearson_global_pos_frac` |
| Pearson global negative fraction | `pearson_global_neg_frac` |
| ***Inter-pocket coupling features*** | |
| Number of other pockets | `n_other_pockets` |
| Inter-pocket coupling mean | `*_to_other_pockets_mean` |
| Inter-pocket coupling max | `*_to_other_pockets_max` |
| Inter-pocket coupling min | `*_to_other_pockets_min` |
| Inter-pocket coupling std | `*_to_other_pockets_std` |
| Max residue-pair coupling | `*_max_pocket_pair_corr` |
| Number of highly coupled pockets | `*_n_high_corr_pockets` |
| Pearson inter-pocket absolute mean | `pearson_to_other_pockets_abs_mean` |
| Pearson inter-pocket absolute max | `pearson_to_other_pockets_abs_max` |
| Pearson inter-pocket absolute std | `pearson_to_other_pockets_abs_std` |
| Pearson inter-pocket positive fraction | `pearson_to_other_pockets_pos_frac` |
| Pearson inter-pocket negative fraction | `pearson_to_other_pockets_neg_frac` |
| Max absolute residue-pair coupling | `pearson_max_pocket_pair_abs_corr` |

Table S2: Benchmarking set (D24)

| PDB ID | Modulator ID | Chain | Modulator Residue No. |
|---|---|---|---|
| 1MC0 | PCG | A | 160 |
| 4OYA | 1VE | A | 501 |
| 2CDQ | SAM | A | 1500 |
| 1WQW | BT5 | A | 1301 |
| 5J94 | 1XF | A | 301 |
| 4PCU | SAM | A | 603 |
| 4NHV | 2O6 | A | 308 |
| 2XEM | SSV | B | 1145 |
| 4DQW | ATP | A | 503 |
| 4TPT | 35H | A | 701 |
| 4FXY | 0W2 | P | 701 |
| 3ZFZ | 1W8 | A | 1669 |
| 3ZFZ | MUR | B | 1669 |
| 4PPV | PHE | A | 401 |
| 5BTR | STL | A | 702 |
| 2C2B | SAM* | A | 501 |
| 4QPL | V3L | A | 201 |
| 4ZLO | 4PV | A | 601 |
| 5CGC | 51D | A | 4006 |
| 4YW8 | 1WD | A | 708 |
| 5DE1 | 59D | A | 502 |
| 5C4T | 4Y6 | A | 601 |
| 3UVV | 9CR | B | 501 |
| 5DED | 0O2 | A | 302 |

*Allosteric modulator (SAM) is located in chain B of the asymmetric dimer.

Table S3: Comparison of fpocket descriptors between biased and unbiased configurations. Biased: *fpocket* applied directly to the holo structure without prior structure preparation. Unbiased: *fpocket* applied after structure preparation, with the allosteric modulator removed from the chain. For 1MC0, the biased configuration corresponds to Pocket 1 and the unbiased configuration to Pocket 7. For 1F1O, both configurations correspond to Pocket 1.

| **Feature** | **1MC0** | | **3F1O** | |
|---|---|---|---|---|
| | **Biased** | **Unbiased** | **Biased** | **Unbiased** |
| Score | 0.358 | 0.014 | 0.937 | 0.630 |
| Druggability score | 0.069 | 0.037 | 0.860 | 0.839 |
| Number of alpha spheres | 50 | 50 | 85 | 85 |
| Total SASA | 53.627 | 138.792 | 1.208 | 76.956 |
| Polar SASA | 24.644 | 57.881 | 0.000 | 8.616 |
| Apolar SASA | 28.983 | 80.911 | 1.208 | 68.340 |
| Volume | 446.241 | 454.061 | 405.616 | 408.810 |
| Mean local hydrophobic density | 20.273 | 20.273 | 38.453 | 38.453 |
| Mean alpha sphere radius | 3.852 | 3.852 | 3.768 | 3.768 |
| Mean alpha sphere solvent accessibility | 0.424 | 0.424 | 0.470 | 0.470 |
| Apolar alpha sphere proportion | 0.440 | 0.440 | 0.624 | 0.624 |
| Hydrophobicity score | 46.200 | 46.200 | 49.682 | 49.682 |
| Volume score | 4.467 | 4.467 | 4.182 | 4.182 |
| Polarity score | 7 | 7 | 9 | 9 |
| Charge score | -3 | -3 | 2 | 2 |
| Proportion of polar atoms | 31.429 | 31.429 | 31.250 | 31.250 |
| Alpha sphere density | 4.382 | 4.382 | 4.958 | 4.958 |
| Center of mass–alpha sphere max distance | 11.266 | 11.266 | 10.556 | 10.556 |
| Flexibility | 0.232 | 0.232 | 0.092 | 0.092 |

Table S4: Pairwise statistical comparison of methods on the unbiased dataset (paired $t$-test across 50 matched splits; seeds sorted by filename before pairing). All distributions passed the Shapiro–Wilk normality test ($p > 0.05$). Cohen's $d$ is computed on paired differences; effect-size thresholds: $|d| < 0.2$ negligible, 0.2–0.5 small, 0.5–0.8 medium. Significance: $^{*}p < 0.05$; $^{**}p < 0.01$; $^{***}p < 0.001$; ns $p \geq 0.05$.

| **Metric** | **Comparison** | $t$ | $p$ | $\|d\|$ | **Effect** |
|---|---|---|---|---|---|
| F1 | Baseline vs AlloDyn (MD) | −3.17 | 0.0026** | 0.45 | small |
| | Baseline vs AlloDyn (AF) | −2.72 | 0.0090** | 0.38 | small |
| | AlloDyn (MD) vs AlloDyn (AF) | −0.02 | 0.9817 ns | 0.00 | negligible |
| MCC | Baseline vs AlloDyn (MD) | −3.21 | 0.0023** | 0.45 | small |
| | Baseline vs AlloDyn (AF) | −2.76 | 0.0080** | 0.39 | small |
| | AlloDyn (MD) vs AlloDyn (AF) | −0.03 | 0.9794 ns | 0.00 | negligible |
| AUC PR | Baseline vs AlloDyn (MD) | −3.94 | 0.0003*** | 0.56 | medium |
| | Baseline vs AlloDyn (AF) | −2.03 | 0.0476* | 0.29 | small |
| | AlloDyn (MD) vs AlloDyn (AF) | 1.51 | 0.1369 ns | 0.21 | small |
| AUC ROC | Baseline vs AlloDyn (MD) | −2.44 | 0.0182* | 0.35 | small |
| | Baseline vs AlloDyn (AF) | −3.69 | 0.0006*** | 0.52 | medium |
| | AlloDyn (MD) vs AlloDyn (AF) | −2.11 | 0.0396* | 0.30 | small |

9 # Supplementary Figures

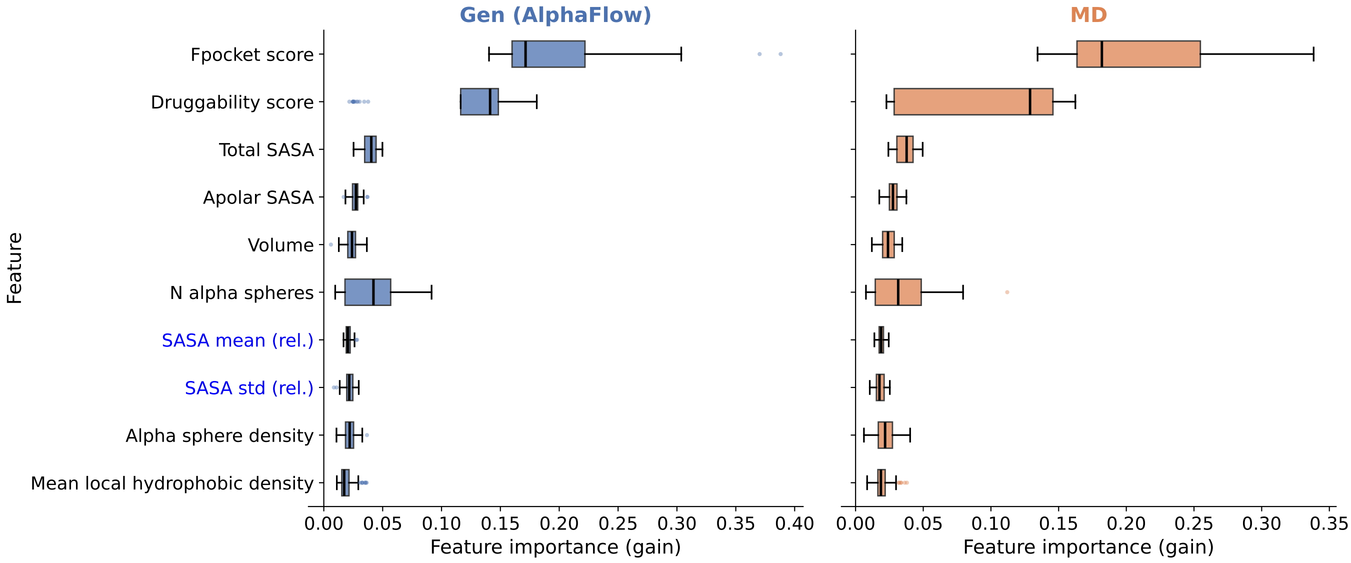


Figure S1: Feature importance of the top-10 fpocket descriptors for biased XGBoost models trained on generated (AlphaFlow) and MD conformations. Importance values (gain, normalised to sum to 1) are shown as horizontal box plots across 50 random train/test splits; features are ordered by mean rank across splits, with the most important descriptor at the top. Pocket score, Druggability score, and Total SASA consistently dominate both models, together accounting for the majority of the total gain. The prominence of these three features is a direct consequence of the data leakage introduced by the biased preprocessing: the presence of the allosteric modulator during *fpocket* feature calculation artificially reduces SASA and, through the regression formulas underlying the score and druggability score, inflates both metrics for allosteric pockets. Feature names shown in blue correspond to dynamic (non-fpocket) descriptors.

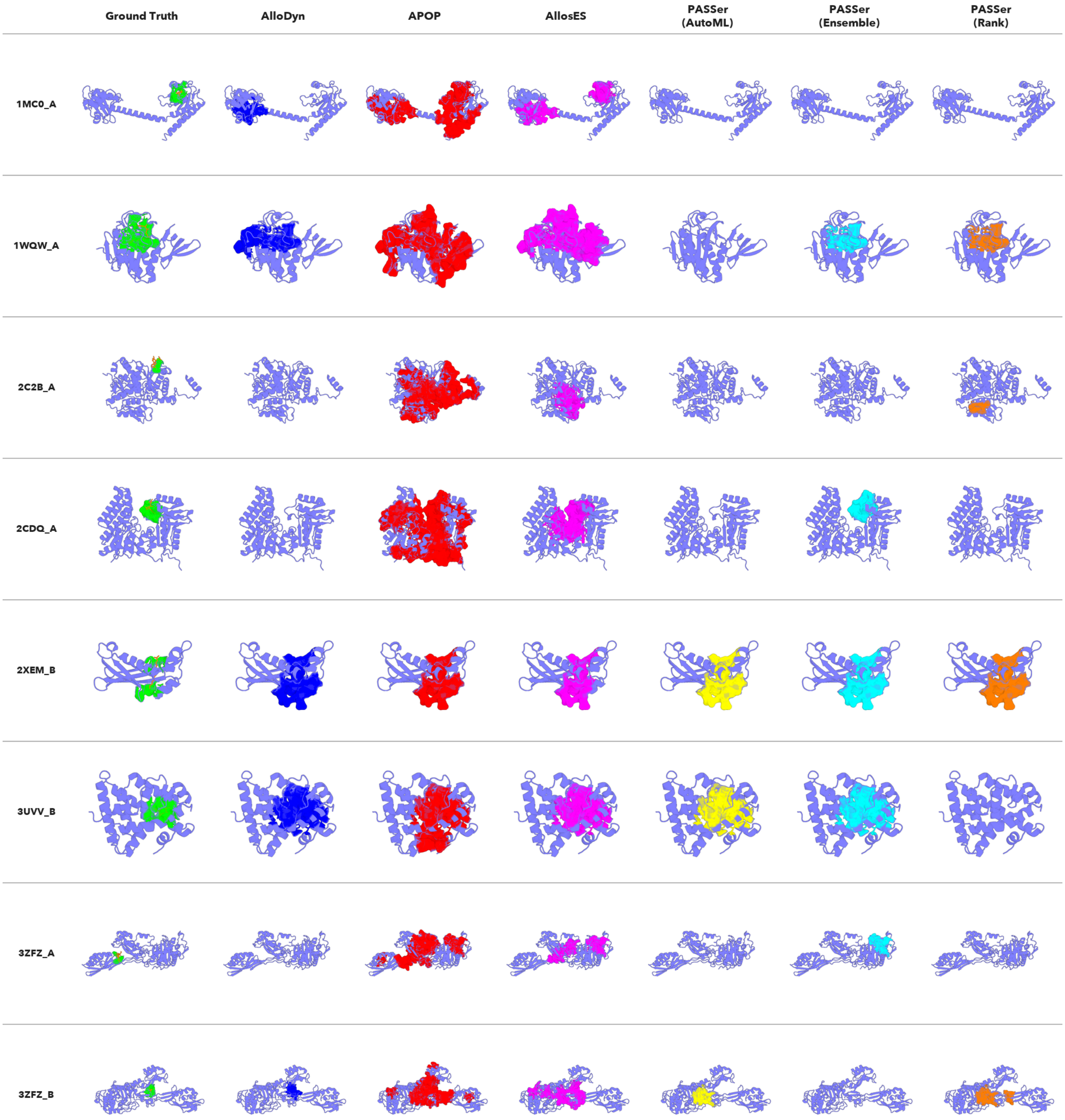


Figure S2: **Predicted allosteric pockets on the D24 benchmark (part 1 of 3).** Each row corresponds to one D24 entry (PDB ID indicated on the left). Columns show the ground truth allosteric site (green, (green, residues within 5 Å of the modulator) and the pockets predicted as positive (allosteric) by each method: AlloDyn (blue), APOP (red), AllosES (magenta), PASSer-AutoML (yellow), PASSer-Ensemble (cyan), and PASSer-Rank (orange). Pockets are displayed as colored surfaces on the protein structure (light blue ribbon). Entries where no pocket is shown indicate that the method failed to return a prediction for that structure.

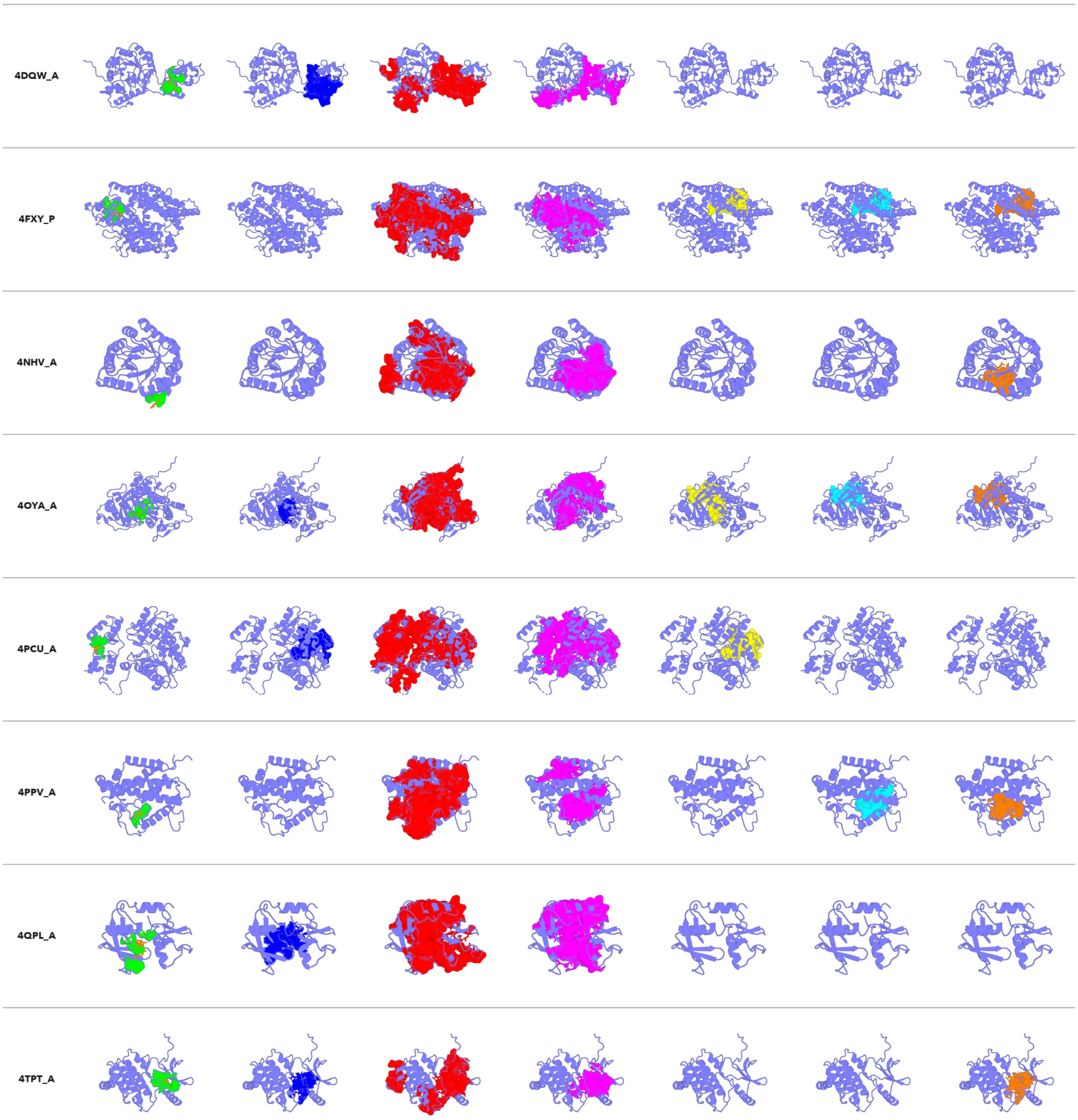


Figure S3: **Predicted allosteric pockets on the D24 benchmark (part 2 of 3).** Continued from Fig. S2.

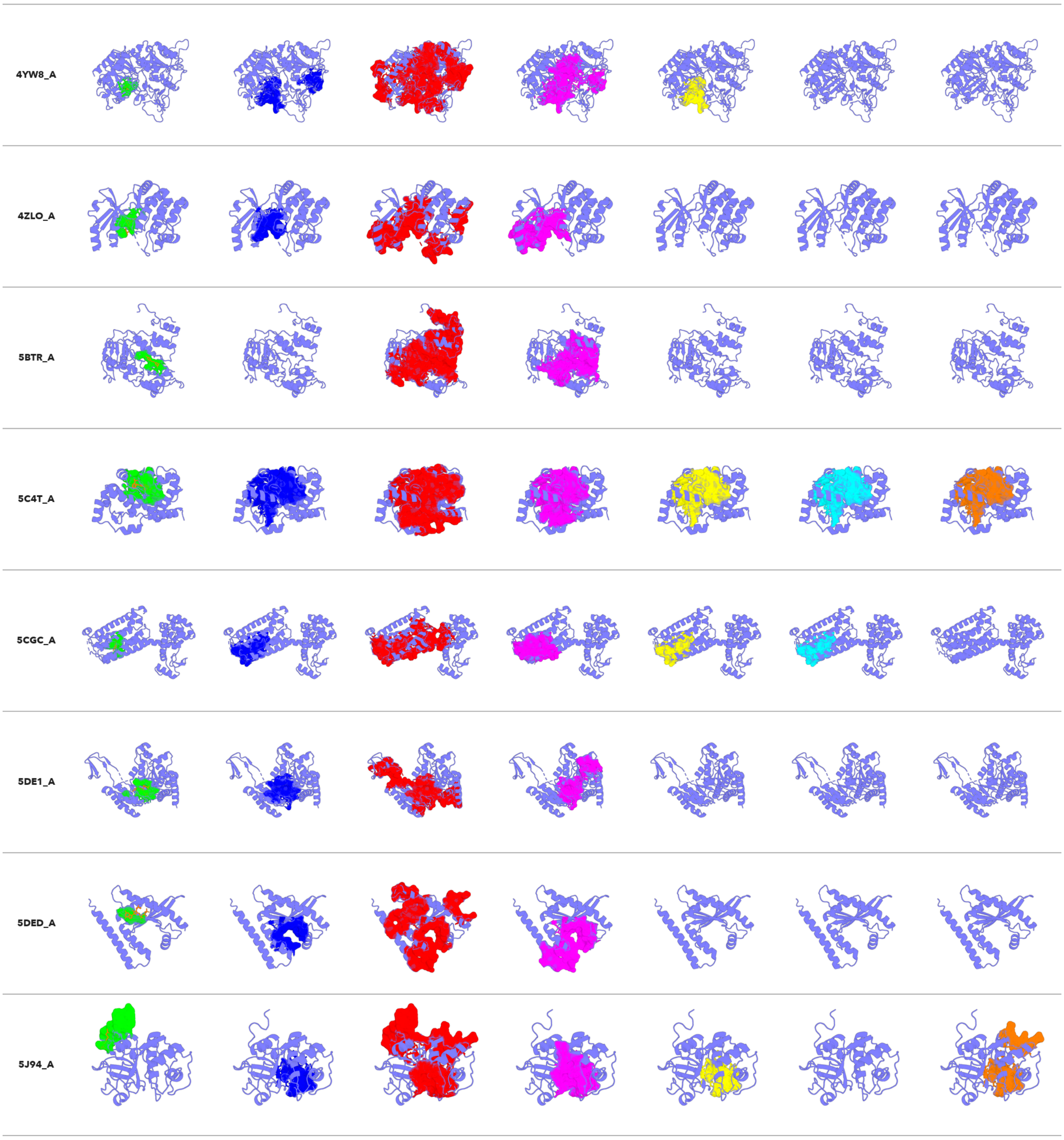


Figure S4: **Predicted allosteric pockets on the D24 benchmark (part 3 of 3).** Continued from Fig. S3.
